# Invariances in a combinatorial olfactory receptor code

**DOI:** 10.1101/208538

**Authors:** Guangwei Si, Jessleen K. Kanwal, Yu Hu, Christopher J. Tabone, Jacob Baron, Matthew Berck, Gaetan Vignoud, Aravinthan D.T. Samuel

## Abstract

Animals can identify an odorant type across a wide range of concentrations, as well as detect changes in concentration for individual odorant type. How olfactory representations are structured to support these functions remains poorly understood. Here, we studied how a full complement of ORNs in the *Drosophila* larva encodes a broad input space of odorant types and concentrations. We find that dose-response relationships across odorants and ORN types follow the Hill function with shared cooperativity but different activation thresholds. These activation thresholds are drawn from a power law statistical distribution. A fixed activation function and power law distribution of activation thresholds underlie invariances in the encoding of odorant identity and intensity. Moreover, we find similar temporal response filters of ORNs across odorant types and concentrations. Such uniformity in the temporal filter may allow identity invariant coding in fluctuating or turbulent odor environments. Common patterns in ligand-receptor binding and sensory transduction across olfactory receptors may give rise to these observed invariances in the olfactory combinatorial code. Invariant patterns in the activity responses of individual ORNs and the ORN ensemble may simplify decoding by downstream circuits.

## Introduction

The abilities to identify odorants across a wide range of concentrations and detect changes in odorant concentration are essential for olfactory perception and behavior. Olfactory systems use combinatorial codes to encode large numbers of odors with smaller numbers of olfactory receptor neurons (ORNs) (Malnic et al., 1999). Each ORN typically expresses one of a large repertoire of olfactory receptors (Ors) (Buck and Axel, 1991). A single Or can be activated by many different odorants, and a single odorant can activate many different Ors (Friedrich and Korsching, 1997). Different odorants can be discriminated by distinct activity patterns across an ensemble of olfactory neurons (Hallem and Carlson, 2006; Kreher et al., 2005; Nara et al., 2011). The olfactory code also conveys information about odorant intensity as higher odorant concentrations tend to activate more ORNs (Kajiya et al., 2001; Wang et al., 2003). Different odorants may also evoke different temporal patterns in neuronal activity, augmenting information coding using time (Friedrich and Laurent, 2001; Laurent et al., 2001; Junek et al., 2010; Smear et al., 2011).

Recent studies have uncovered coding properties at the single cell and population levels that may allow for scale-invariant representation of olfactory information such as odorant type and intensity. At the individual ORN level, ORN responses to temporal patterns in odorant presentation may be converted into predictable activity patterns by stereotyped filters (Nagel and Wilson, 2011; Martelli et al., 2013). At the population level, inputs to the olfactory bulb may encode odorants in concentration invariant spatial representations (Wachowiak et al., 2002; Cleland et al., 2007). At the statistical level, the firing rates of *Drosophila* ORNs appear to be drawn from an odor invariant probability distribution (Stevens, 2016). However, a quantitative characterization of such invariances in olfactory representation by a complete ORN ensemble is still missing.

In this study, we characterized the ORN ensemble of the *Drosophila* larva to a panel of odorant types and concentrations that spanned the selectivity of all olfactory sensory neurons. The *Drosophila* larva offers the advantage of numerical simplicity for dissecting an olfactory circuit that shares glomerular organization with adult insects and vertebrates (Vosshall and Stocker, 2007; Su et al., 2009). We find that ORN-odorant pairs share the same activation function: ORN activity increases with concentration along the same Hill curve for any odorant type but with odorant-specific thresholds. We find that the statistical distribution of these ORN sensitivities to odorants across olfactory space follows a power-law. Furthermore, ORNs share a stereotyped temporal filter shape such that ensemble level responses may be concentration-invariant in a fluctuating environment. Our systems-level characterization of an entire olfactory periphery across a wide range of odorant types and concentrations has revealed individual and ensemble level ORN patterns that allow for invariant representation of olfactory information, with significance for downstream processing.

### A microfluidic setup for *in vivo* calcium imaging of larval ORNs

Small size and optical transparency make the larva’s olfactory system - like that of *C. elegans-* suitable for *in vivo* multineuronal calcium imaging with precise and flexible microfluidic control of olfactory inputs (Chronis et al., 2007). We developed a microfluidic device for an intact, un-anesthetized larva with up to 16 fluid delivery channels, allowing us to image olfactory processing in single animals exposed to a broad input space (**Fig 1A-D**, **Supp Fig 1**). Fluid delivery allows for precise control of odorant concentration, timing between stimulus delivery, and stimulus waveform (Andersson et al., 2012). Furthermore, with the microfluidics setup we can record from ensembles of olfactory neurons with single cell resolution while delivering inputs that span odorant types and concentrations. Calcium imaging and genetic labeling allow us to record the activity of any individual ORN alone or the activity of all ORNs simultaneously, by expressing the calcium indicator GCaMP6m (Chen et al., 2013) under the control of either a specific ORN *Gal4* driver or the *Orco-Gal4* driver, respectively.

**Figure 1.**
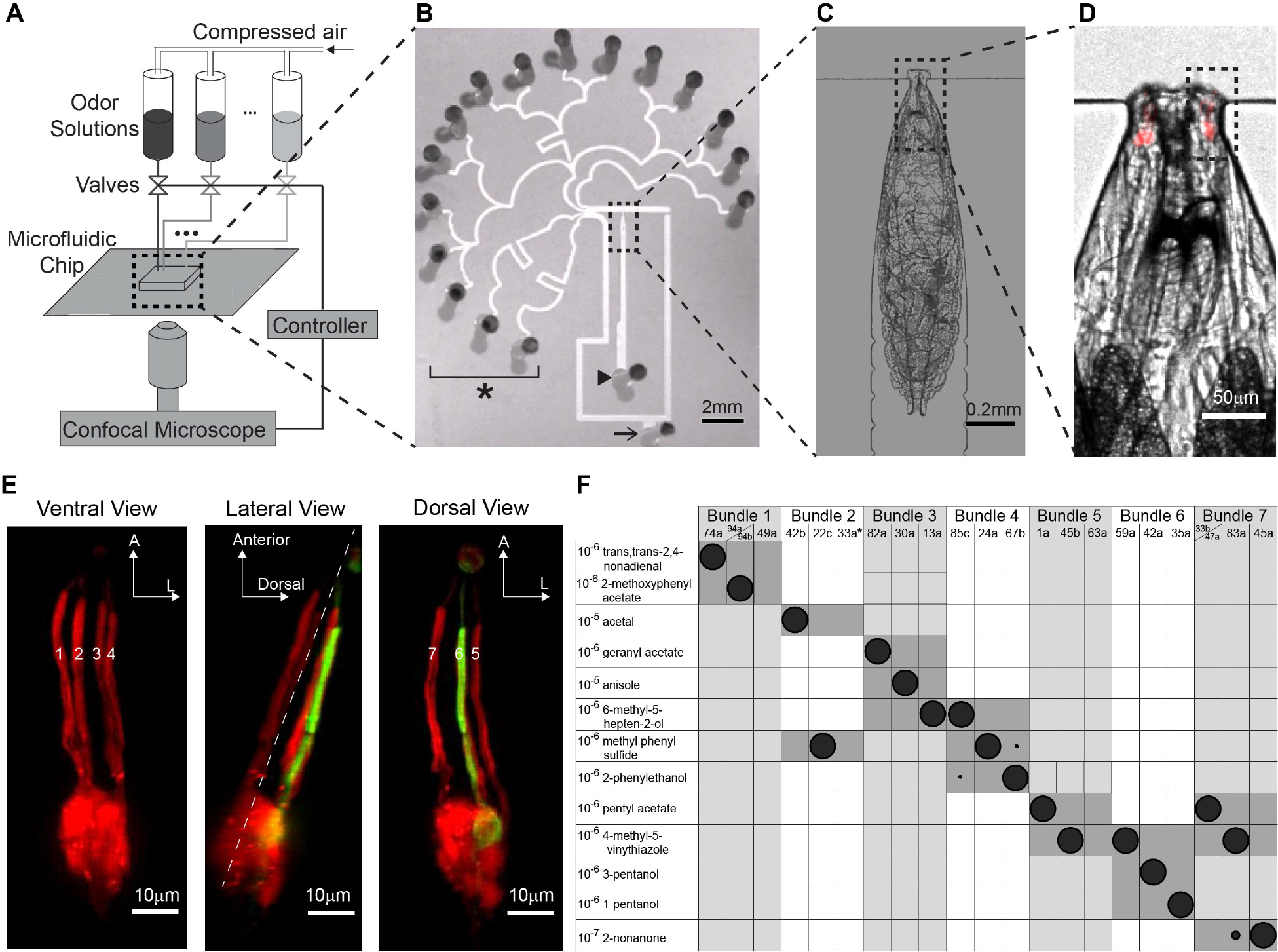
Anatomical and functional identification of individual ORNs within the ORN population. A. Schematic of the setup for confocal imaging of a larva in a microfluidic device during odorant delivery. B. 16 channel microfluidic device (* indicates stimulus delivery channels, arrowhead marks larva loading inlet, arrow marks fluid outlet). C and D. Zoomed-in view of an immobilized larva in the loading channel. Red indicates RFP labeling of ORN dendrites and cell bodies. E. Larval ORN sensory dendrites are organized into seven parallel bundles (numbered). All ORNs shown in red, Or35-ORN shown in green, using *Or35a>GFP; Orco>RFP* genotype. Dashed line in lateral view marks separation between ventral and dorsal bundles. F. Each of 13 odorants at low concentrations primarily activates only one ORN within each bundle. Size of shaded circles indicates normalized neural activity (ΔF/F) of the specified ORN to an odorant. * indicates that location of Or33a-ORN was inferred from vacancy in bundle 2 (**Supp Fig 2**).

### Anatomical and functional identification of individual ORNs

The larva has 21 ORNs located in each bilaterally symmetric dorsal organ ganglion (DOG). The layout of ORN dendrites aids in segmenting and identifying all cells during multineuronal calcium imaging. The 21 ORN sensory dendrites are organized into seven parallel bundles, each containing three sensory dendrites, that project from an ORN soma to the dorsal organ, a perforated dome on the animal’s head (Singh and Singh, 1984). When a larva is immobilized in the microfluidic device, four ventral and three dorsal dendritic bundles are easily distinguished (**Fig 1E**). We mapped individual ORNs to each bundle by expressing RFP in all ORNs and GFP in a selected ORN using a cell specific Gal4 driver (**Supp Fig 2**). We found that the three ORN dendrites located in each bundle were stereotyped (confirmed in n ≥ 5 animals for each cell type). Thus, by following the activation of any cell body in the DOG to its corresponding dendritic bundle, its possible identity is narrowed to one of three ORNs.

To further aid in the identification of individual ORNs, we used a set of odorants, termed private odorants, that activate single ORNs at low concentrations. Mathew et al. (2013) assembled a panel of 18 private odorants for each larval ORN by expressing a single functional larval olfactory receptor (Or) in a mutant adult ORN devoid of the endogenous Ors, and recorded its electrical activity in response to olfactory cues. We delivered these private odorants to larvae in our microfluidic setup and found that 18 of the 21 ORNs, in each DOG, are responsive to these odorants: none of the private odorants in the panel activate the Or33a or Or63a ORNs and the Or49a ORN is only activated by a wasp pheromone (Ebrahim et al., 2015). We found that 13 of the private odorants are sufficient to identify all ORNs when examined in conjunction with dendritic bundle location (**Fig 1F**). Together, the anatomical map and functional responses to this subset of private odorants provides a comprehensive means of identifying and segmenting the ORNs responsive to any olfactory input during multineuronal imaging.

### ORN ensemble responses across odorant identities and intensities

The panel of 18 private odorants provides a maximally decorrelated set of stimuli that spans the larval olfactory system. To characterize the olfactory representation of these stimuli, we exposed larvae to all 18 private odorants across the concentration range of olfactory sensitivity. We measured the response amplitude of every cell to step stimuli across five orders of magnitude in concentration, from 10^−8^ dilution (where all private odorants were at or below threshold of ORN detection) to 10^−4^ dilution (where many ORNs had reached saturation). We used five second step pulses interleaved with 20-60 seconds of water, a protocol that allowed us to measure peak responses and allowed for full recovery of neural activity (**Supp Fig 3**).

We verified that all private odorants were highly selective for their target ORNs at low concentrations, with activity expanding to additional ORNs at higher concentrations. For example, 1-pentanol was identified as a private odorant for the Or35a-expressing ORN. At 10^−7^ dilution, 1-pentanol slightly evoked activity specifically in the Or35a-ORN. Higher concentrations of 1-pentanol gradually saturated the Or35a-ORN, while also activating four other ORNs expressing either Or67b, Or85c, O13a, or Or1a (**Movie 1**). Interestingly, each additional ORN recruited by 1-pentanol corresponded to a private odorant that is also a long chain alcohol (Mathew et al., 2013). We next examined the ensemble-wide dose-response curves for these additional private alcohol odorants. Low concentrations of each private alcohol specifically activated its target ORN. Higher concentrations reliably activated the Or35a-ORN, Or13a-ORN, Or67b-ORN, and Or85c-ORNs to varying degrees (**Supp Fig 4**). Furthermore, we performed the dose-response analysis across the entire ORN ensemble for all 18 private odorants (**Fig. 2A**). We found a similar pattern of overlapping activation for ORNs sharing an odorant with a similar molecular structure. Thus, as odorant concentration increases, a family of molecules with similar structure will cross-activate the subgroup of ORNs that are particularly selective for molecules within the same family.

**Figure 2.**
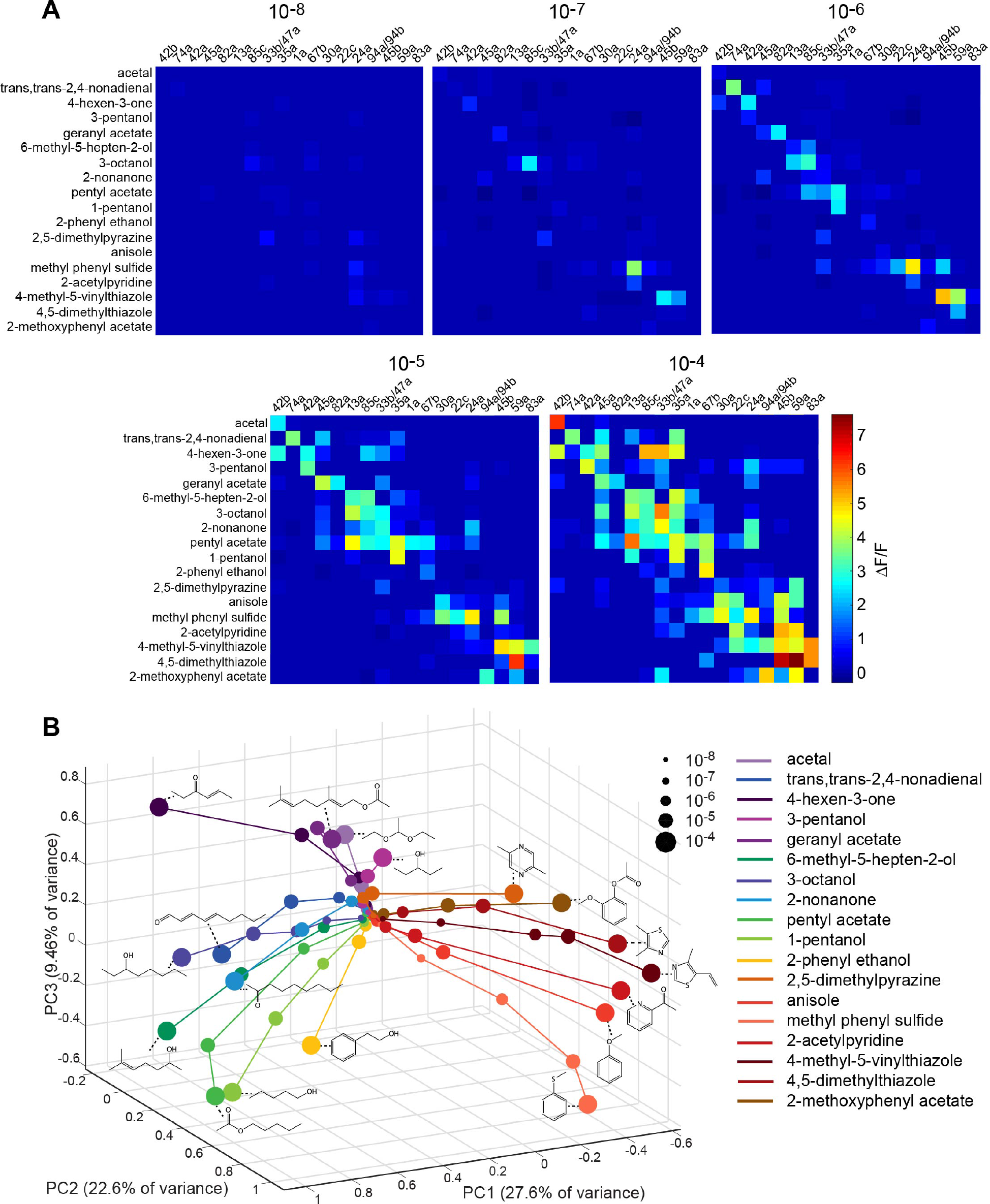
ORN population responses to different odorants and concentrations. A. Averaged peak responses of 18 ORNs to a panel of 18 odorants, each delivered at five concentrations (n ≥ 5 for each odorant type and concentration; odorant pulse = 5 s). B. ORN population responses visualized in PCA space. Each dot represents the projection of ORN population activity onto the first three principal components. Size and color of dots correspond to odorant concentration and type, respectively. Dots from the same odorant are linked and the molecular structure of the odorant is shown adjacent to each trajectory. Aromatic versus aliphatic odorants cluster in separate regions of PCA space.

As in other animals, the olfactory code changes with increasing odorant intensity (Malnic et al., 1999), but with a pattern of ORN recruitment that is correlated with molecular selectivity. To discern this pattern, we used principal component analysis (PCA) of the responses of all ORNs measured against all private odorants across all concentrations. We visualized the data by projecting the ORN activity responses in the space of the first three principal components (PCs) (**Fig 2B**), which account for 60% of the variance in the data (**Supp Fig 5A**). At the lowest concentrations, olfactory representations at or below the detection threshold across odorants were tightly clustered at a central point in the PCA space. At higher concentrations, olfactory representations diverged, increasing distance monotonically from the central point (**Fig 2B**, **Supp Fig 5B**). Interestingly, the trajectory of each odorant tended to follow its own direction in PCA space. This pattern is particularly clear for aliphatic and aromatic odorants. Aliphatic odorants with long carbon chains form trajectories projecting in a similar direction of PCA space, since higher concentrations of these odorants tend to selectively recruit the other ORNs with aliphatic private odorants. The same was true for aromatic odorants and the corresponding group of ORNs with private odorants of this type (**Fig 2B**). The vectors corresponding to structurally similar molecules were separated by small angles (**Fig 2B**, **Supp Fig 5C**). Thus, visualization of ORN responses in PCA space reveals structure in the ensemble representation of odorant identity over a large range of intensities. The population wide response maintains a fixed direction in the representation of each odorant as concentration rises.

### Dose-response curves share the same steepness but vary in threshold concentrations

We uncovered additional invariant structure when we analyzed the dose-response relationship of individual odorant-ORN pairs. We found that the subset of all pairs that reached saturation (n= 21 of 324 pairs) were well described by a Hill function:

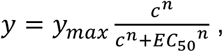

where *y*_*max*_ is the maximum response amplitude measured by the calcium indicator, *c* is the odorant concentration, *n* is the Hill coefficient or steepness of the linear portion of the curve, and*EC*_50_ is the half-maximal effective concentration. The Hill function canonically describes binding affinities in ligand-receptor interactions such as that between odorants and olfactory receptors. Here, we find that the Hill equation describes a common concentration dependent nonlinearity in each dose-response relationship. After normalizing each dose-response curve by *y*_*max*_ and aligning by the *EC*_50_, all 21 dose-response curves collapsed onto a single Hill function with *n* = 1. 5 ± 0.1 (**Fig 3A**). This common Hill coefficient suggests a similar degree of cooperativity in odorant binding and signal transduction across the ORN repertoire. Assuming the same cooperativity applies to the other odorant-ORN pairs, we estimated the *EC*_50_ value for all remaining pairs. The complete *EC*_50_ matrix reveals the distribution of sensitivities across the ORN ensemble to each odorant (**Fig 3B**).

**Figure 3.**
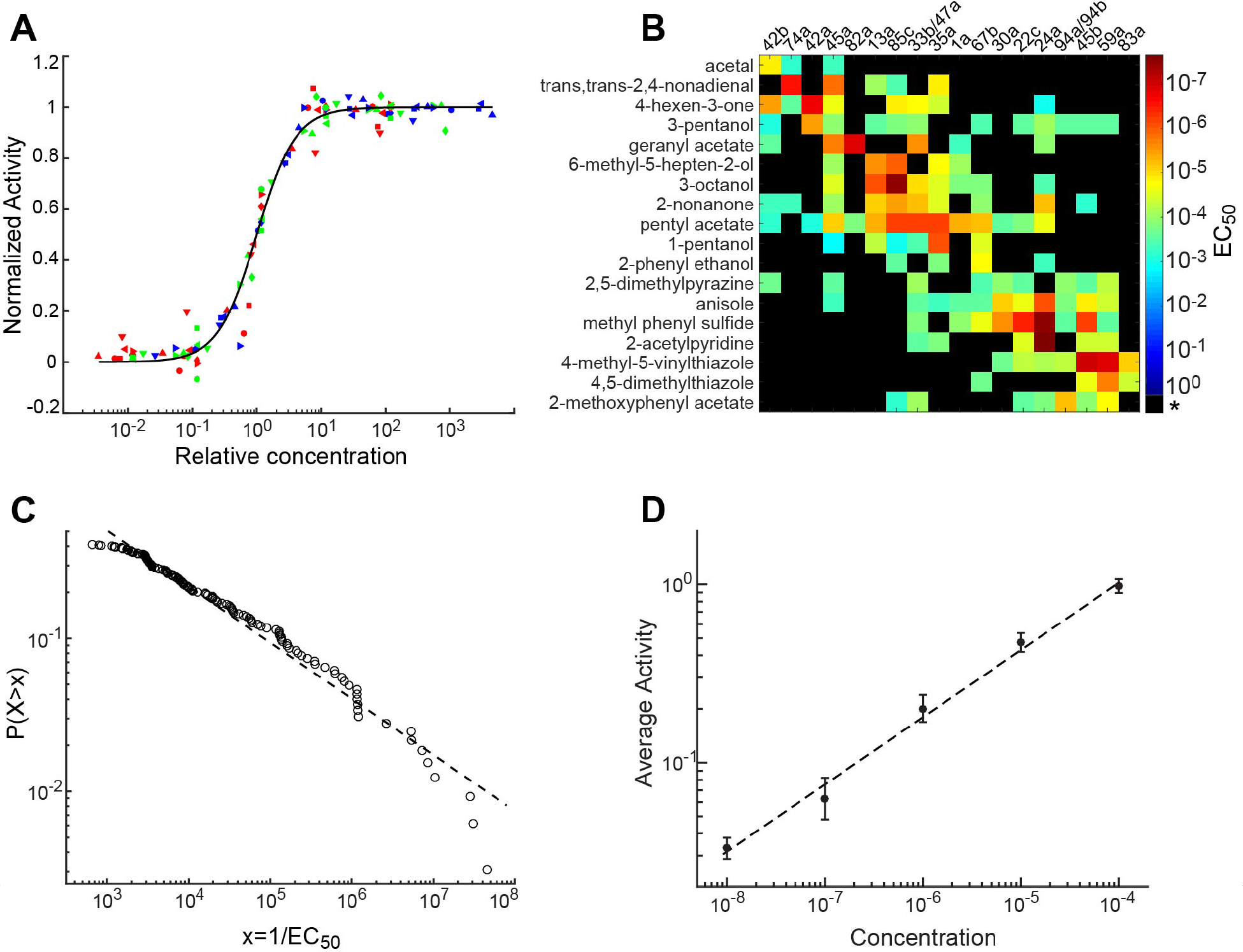
Scaling laws for individual and ensemble ORN activity. A. Normalized ORN responses for various odorant-ORN pairs across relative odorant concentration (actual concentration divided by *EC*_50_). Individual curves for plotted odorant-ORN pairs collapse onto a single curve described by a Hill equation with a shared Hill coefficient of 1.45. Black line indicates the fitted Hill equation, different colored and shaped points represent data from unique odorant-ORN pairs. B. Matrix of *EC*_50_ values fit to dose-response data from each odorant-ORN pair (* for black squares indicates that odorant-ORN pair had no response within the tested concentration range). C. Log-log plot of the cumulative distribution function of 1/*EC*_50_ values. The dashed line is a linear fit to the data and has a slope of −0.35. D. Log-log plot of average neuron activity across all odorant-ORN pairs for each concentration. The error bars represent the standard error. Least-squares fit line has a slope of 0.38±0.06 (*R*^2^ = 0.99).

A simple coding scheme emerges. A common Hill function, with the *EC*_50_ value as the only free parameter, describes the dose-response relationship for any odorant-ORN interaction. This model, using the complete matrix of estimated *EC*_50_ values, accounts for 98% of the variance in the original dataset (**Supp Fig 6A**). For each odorant, the vector of *EC*_50_ values (a row in the matrix in **Fig 3B**) specifies the identity and threshold of each activated ORN with increasing odorant concentration. A corollary of having a unique *EC*_50_ vector for each odorant is having a unique direction for the trajectory of population responses across concentrations (**Fig 2B**).

To study structure in the distribution of ORN sensitivities, we applied PCA to the matrix of ln(1/*EC*_50_) (see **Methods**). We found that the first principal component (PC) explains a significant portion of the variance (**Supp Fig 6B**). We projected the vector of ln(1/*EC*_50_) values associated with each private odorant onto this first PC, and found that this projection strongly correlated with aromaticity index (**Supp Fig 6C**), one of the major quantitative metrics of odorant molecular structure that has been linked to olfactory discrimination across animals (Haddad et al., 2008). This observation explains why the trajectories of aromatic and aliphatic odorant representations point in opposite directions in **Fig. 2B**.

### Power law distribution of ORN ensemble sensitivities

Next, we examined the properties of the *EC*_50_ values themselves. We extracted all measured elements from the *EC*_50_ matrix and constructed a cumulative density function (**Fig. 3C**). The data closely follows a line in the log-log plot, indicating a power law: 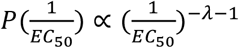, λ = 0.35.

A power law distribution of olfactory sensitivities means that a relative change of concentration will trigger the same mean relative change in the number of activated ORNs, irrespective of odorant type. A power law distribution of olfactory sensitivities, together with a common Hill function, should give rise to ensemble-wide activity that follows a power law relationship with respect to concentration and has an exponent *λ* (See **Methods**). We confirmed this prediction in our experimental data (**Fig 3D**). The mean activity of the olfactory ensemble grows with odorant concentration following a power law with an exponent of 0.38±0.06, which is close to the 0.35 exponent found from fitting the *EC*_50_ matrix (**Fig 3C**). Thus, on average, activity expands across the ORN ensemble at the same rate with increasing relative concentration, irrespective of odorant type (as shown in **Fig. 2**).

### ORN-odorant responses share similar temporal characteristics

An additional challenge to olfactory coding of a wide variety of odorant types across concentrations arises from complex temporal dynamics due to physical fluctuations, such as turbulence or convection, in the stimulus itself. To examine how such fluctuations affect ORN responses, we compared the conversion of temporal patterns of olfactory input for different odorant-ORN pairs across odorant intensities. To do this, we used reverse-correlation analysis, subjecting larvae to “white noise” olfactory input by stochastically switching between odorant and water delivery and seeking the temporal filter that best maps olfactory inputs into calcium dynamics (Geffen et al., 2009; Kato et al., 2014). We found that random olfactory input could evoke fluctuating calcium activity in an ORN, and repeated presentation of the same input pattern would evoke consistent responses from trial to trial (**Supp Fig 7A**). The systematic conversion of the stimulus to response waveform is well characterized by a linear-nonlinear (LN) model. A linear transfer function estimates the relative weight of each time point in stimulus history to determine the time-varying response amplitude (**Supp Fig 7B**). The convolution of the linear transfer function with stimulus history is then passed through a static nonlinearity to correct for saturation (**Supp Fig 7C**). We verified the LN model by predicting the response to a novel random input using a filter calculated from different random inputs (**Supp Fig 7D**).

We measured the linear transfer function for 3-octanol, the private odorant for the Or85c-ORN, across the concentration range used to characterize the *EC*_50_ matrix. At the lowest concentrations of 3-octanol, a filter describing ORN activity only emerges for the Or85c-ORN (**Fig 4A**). At higher concentrations, filters begin to emerge for additional ORNs. These filters for each ORN, when normalized for response amplitude, were virtually identical in their temporal response profiles as single lobed functions with similar peak and decay times (**Fig 4B**, **Supp Fig 8B-C**). The shapes of the filters for different odorants activating the same ORN are also virtually indistinguishable, on the order of ~100 ms (**Fig 4C**, **Supp Fig 8A-C**). This result is constrained by the calcium indicator, which has a time constant associated with calcium binding to GCaMP6m, making it difficult to resolve differences in ORN temporal filters on a shorter time scale. Nonetheless, recent electrophysiological measurements of odorant-evoked activity in the ORNs of adult *Drosophila*, under the same LN model, also report remarkable similarity in the temporal pattern of filters across odorant-receptor pairs, within ~10-20 ms (Martelli et al., 2013).

**Figure 4.**
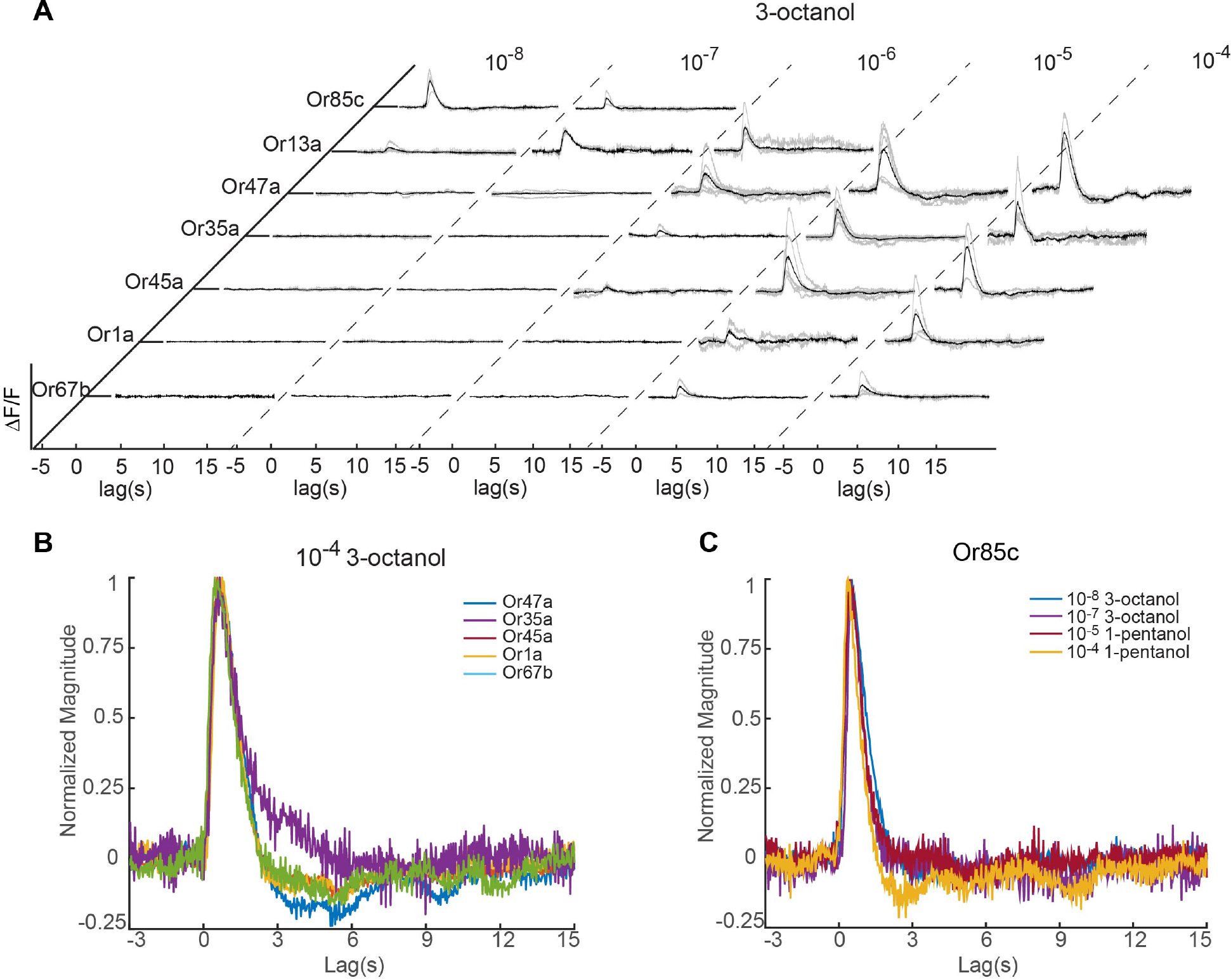
Temporal filters of ORN response. A. Linear filters of 7 ORNs responding to 3-octanol across five concentrations. Black curve indicates the averaged filter from data across multiple animals (individual filters shown in gray). B and C. Comparison of filter waveforms for the same odorant (10^-4^ dilution of 3-octanol) activating different ORNs (B), and the same ORN (Or85c-ORN) responding to different odorants and concentrations (C). All filters were normalized by their peak amplitude.

A common temporal filter across ORNs could simplify the olfactory code in an environment with fluctuating odorant concentrations. A constant filter in conjunction with uniform scaling of ORN activity over concentration could allow the ensemble of responsive neurons to maintain the same relative amplitudes of activation over time. These relative amplitudes would be correlated with the ORN ensemble *EC*_50_ values for any odorant, regardless of whether an animal is in a static or fluctuating odorant environment.

## Discussion

Previous efforts at a systems-level characterization of ORNs necessarily focused analysis on particular cell types, odorants, or odorant concentrations (Hallem and Carlson, 2006; Mathew et al., 2013; Nagel and Wilson, 2013; Martelli et al., 2013; Asahina et al., 2009). The small size of the *Drosophila* larva, combined with multineuronal imaging and new microfluidic tools, has allowed us to characterize the responses of a complete ORN ensemble to a panel of odorant types and concentrations that spans the selectivity of olfactory sensory neurons. This broad characterization has uncovered regular patterns in the response of individual ORNs and of the ORN ensemble. First, each ORN response to odorants exhibits the same activation function shape with variant sensitivity levels. Furthermore, consistent temporal filters that convert different stimulus waveforms into ORN calcium activity patterns will make the relative activities of different ORNs robust despite fluctuating inputs. Second, the ORN ensemble across all tested odorants exhibits a constant rate of increase in activity with increasing concentration. Underlying this effect, we have identified a power-law distribution in the sensitivities of odorant-ORN interactions. The power law distribution may allow downstream circuits to estimate the relative concentrations of any odorant by the relative extent of ORN activity using the same quantitative relationship for any odorant. Invariant quantitative patterns in single and ensemble-level ORN activities could allow shared mechanisms to extract olfactory features, even in fluctuating environments, across olfactory space.

Different neurons are required to sense odorants in different regimes of odorant concentration needed for long-range chemotaxis in the *Drosophila* larva (Asahina et al., 2009). Encoding a broad concentration range requires a distribution of ORNs with varying sensitivities. Our analysis reveals that olfactory sensitivities follow an invariant statistical distribution across odorants and ORN types. This power law distribution implies a fixed ratio between the relative change in ORN ensemble activity for a fixed change in odorant concentration. Detection of relative change in stimulus intensities has been observed in psychophysical studies of diverse sensory modalities. A notable example is Stevens’s Law in human psychophysics, where response magnitudes have been shown to scale with the logarithm of stimulus intensities across sensory modalities including olfaction (Stevens, 1957). Our results reveal that a phenomenon analogous to Stevens’s Law can be attributed to the olfactory sensory periphery itself, a direct outcome of the statistical distribution of response sensitivities across an ensemble.

A combinatorial olfactory code will arise from a distribution of ORN sensitivities to different odorant molecules. Changes in concentration necessarily lead to changes in the combinatorial code, but with correlated changes depending on the odorant. The basis of this correlation is a unique vector of sensitivities across ORNs for each odorant. This constraint allows each odorant’s identity to be coded in a concentration-independent manner as a direction in an olfactory coding space. It has been suggested that extracting relative glomerulus activity across odorant concentrations may allow the concentration-invariant coding of different odorant types (Wachowiak et al., 2002; Cleland et al., 2007). For animals that sniff, the change in concentration through inhalation generates a reliable temporal sequence of ORN activity that could represent the vector of ORN sensitivity (Smear et al., 2011). As we have found in *Drosophila*, a common activation function and temporal filter - which may arise from stereotyped receptor cooperativities and shared transduction dynamics among ORNs - would facilitate the decoding that takes place by such mechanisms to extract concentration invariant representations of odorant identity.

To our knowledge, a power law distribution of olfactory sensitivities has not yet been described in any animal. One possibility for the power law in olfactory sensitivity is to match the distributions of odorant concentrations found in natural olfactory environments. Natural odors are mixed by convection and turbulence, physical processes that are rich in power law dynamics (Catrakis and Dimotakis, 1996). Power laws appear in the statistics of other natural stimuli as well. Natural visual scenes exhibit a power law relationship between spectral power and spatial frequency (Field, 1987; Simoncelli and Olshausen, 2001). The loudness of natural sounds across frequencies are distributed by power laws (Theunissen and Elie, 2014). Sensory systems, in general, may adapt the statistical distribution of their sensitivities to their natural environments.

Another possibility is that the observed power law distribution represents an optimization of the olfactory code. It has recently been proposed that the olfactory code maximizes informational entropy (Stevens, 2015). Given the constraint of fixed mean firing rates among ORNs, this model leads to an exponential distribution of ORN firing rates evoked by odorants measured in the adult *Drosophila* antenna (Stevens, 2016). Interestingly, another prediction of this optimization is that the overall activity of the olfactory ensemble should increase as a power law of odor concentration, as we have also experimentally observed in the *Drosophila* larva and, in our case, connected to the statistical distribution of olfactory sensitivities across ORNs.

Finally, the molecular recognition mechanism of olfactory receptors may themselves give rise to a power law distribution in ORN sensitivities. Lancet et al., 1993 proposed a molecular recognition system in which a receptor has multiple selective binding sites. Each binding site contributes in a combinatorial manner to the binding strength between a receptor and molecule. This simple quantitative model generates a power law sensitivity distribution for receptors with random sets of binding sites. The statistics of an olfactory code using such a molecular recognition system would be robust to expansion of the ORN periphery, as occurs with *Drosophila* in which the adult has nearly triple the number of receptor types as that found in the larva. Furthermore, a conserved statistical structure in the olfactory code would allow downstream circuitry to employ similar decoding mechanisms across an animal’s lifetime.

## Acknowledgment

The authors would like to thank the valuable suggestions from Benjamin de Bivort, Dmitri Chklovskii, Cengiz Pehlevan, Marta Zlatic, Betty Hong, and Yuhai Tu. This work was performed in part at the Harvard Center for Nanoscale Systems, a member of the National Nanotechnology Infrastructure Network (NNIN), which is supported by the National Science Foundation under NSF award no. ECS-0335765. This work was also supported by a National Science Foundation Graduate Research Fellowship (DGE1144152), NSF Brain Initiative grant (NSF-IOS-1556388), and grants from the NIH (8DP1GM105383, P01GM103770, F31DC015704). YH acknowledges support from the Swartz Program in Theoretical Neuroscience at Harvard.

## Methods

### Fly stocks

Flies were reared at 22°C under a 12:12 hour light/dark cycle in vials containing conventional yeast agar medium. Adult flies were transferred to a larvae collection cage (Genesee Scientific) containing a grape juice agar plate and a dime-sized amount of fresh yeast paste. Flies could lay eggs on the grape juice agar plate for two days and then the plate was removed for collection of first instar larvae. The following fly lines were used in this study: *UAS-mCherry.NLS; UAS-GCaMP6m, UAS-mCD8::GFP; Orco::RFP, Orco-Gal4* (BL23292), *Or1a-Gal4* (BL9949), *Or13a-Gal4* (BL9945), *Or22c-Gal4* (BL9953), *Or24a-Gal4* (BL9958), *Or30a-Gal4* (BL9960), *Or33b-Gal4* (BL9963), *Or35a-Gal4* (BL9968), *Or42a-Gal4* (BL9970), *Or42b-Gal4* (BL9971), *Or45a-Gal4* (BL9976), *Or45b-Gal4* (BL9977), *Or47a-Gal4* (BL9982), *Or49a-Gal4/Cyo; Dr/TM3* (gift from John Carlson lab), *Or59a-Gal4* (BL9990), *Or63a-Gal4* (BL9992), *Or67b-Gal4* (BL9995), *Or74a-Gal4* (BL23123), *Or82a-Gal4* (BL23125), *Or83a-Gal4* (BL23128), *Or85c-Gal4* (BL23913), *Or94b-Gal4* (BL23916).

### Microfluidic device design, fabrication, and calibration

Odorant stimuli were delivered using a microfluidic device (**Fig 1A**) designed with a 300 μm wide and 70 μm high larva loading channel. The channel tapered to a width of 60 μm in order to immobilize the larva. The tapered end was positioned perpendicular to a stimulus delivery channel to allow for odorant flow past larval ORNs. The device was designed with a “shifting-flow strategy”, similar to that described in Chronis et al, 2007. The 16-channel device included two control channels located at the periphery, 13 odorant channels in the middle, and one water channel to remove odorant residue. Each channel was of equal length to ensure equal resistance. During an experiment, a combination of three channels was always open: the water channel, one of the 13 odorant delivery channels, and one of the control channels. The 13 odorant channels could be sequentially opened to deliver any odorant. Switching between the two control channels directed either water or an odorant to flow past the larva’s ORNs, as demonstrated in **Supp Fig 1**.

Fluorescein dye was used to measure the switching time between water and odorants as well as to verify the spatial odorant profile in the device during stimulus delivery. Our standard air pressure for stimulus delivery was 6 psi, which led to a flow rate of 0.5 mL/min in the microfluidic device. With these conditions, the switching time between water and odorant was ~20 ms (**Supp Fig 1A**).

The microfluidic device pattern was designed using AutoCAD. The design pattern was then transferred onto a silicon wafer using photolithography. The wafer was used to fabricate microfluidic devices using polydimethylsiloxane (PDMS) and following the standard soft lithography approach (Anderson et al, 2000). The resulting PDMS molds were cut and bonded to glass cover slips. Each microfluidic device was used for only a single panel of odorants in order to prevent contamination.

### Odorant delivery setup

Odorants were obtained from Sigma-Aldrich, diluted in deionized (DI) water (Millipore) and stored for no more than 2 days. To prevent contamination, each odorant concentration was stored in a separate glass bottle and delivered through its own syringe and tubing set. Panels of odorants were delivered using a 16-channel pinch valve perfusion system (AutoMate Scientific, Inc.). Each syringe and tubing set contained a 30 mL luer lock glass syringe (VWR) connected to Tygon FEP-lined tubing (Cole-Parmer), which in turn was connected to silicone tubing (AutoMate Scientific. Inc.). The silicone tubing was placed through the pinch valve region of the perfusion system as its flexibility could allow for the passage or blockage of fluid flow to the microfluidics device. The silicone tubing was then connected to PTFE tubing (Cole-Parmer), which was then inserted into the microfluidic device. A microcontroller and custom written Matlab code were used to control the on/off sequence of the valves and to synchronize valve control with the onset of recording in the imaging software (NIS Elements).

During the entire recording, the larva experienced continuous fluid flow, with a flow rate of 0.5mL/min or 0.2m/s. In the dose-response experiments, the stimuli sequences consisted of five seconds of odorant pulses followed by a washout period using water. The duration of odorant pulses was chosen such that ORN responses reached maximum amplitude. The washout time was adjusted to allow for ORN recovery back to baseline activity levels, and thus ensured that measurements of ORN responses were independent of stimulus sequence (**Supp Fig 3** and **Movie 1**). For the white noise experiments, a 1024-step m-sequence of odorant stimulus and water was delivered with a time step of 0.2 s (**Movie 2**).

### Calcium imaging

A first instar larva was loaded into a microfluidic device using a 1 mL syringe filled with 0.1% triton-water solution. Using the syringe, a larva was pushed towards the end of the channel, where the 60 μm wide opening mechanically trapped further larval movement. Each larva was positioned such that its dorsal organ (nose) was exposed to the stimulus delivery channel and its dorsal side (where ORN cell bodies are located) was closest to the objective. Larvae were imaged using an inverted Nikon Ti-e spinning disk confocal microscope with a 60X water immersion objective (NA 1.2). A charged-coupled device (CCD) microscope camera (Andor iXon EMCCD) captured images at 30 frames/sec. ORN cell bodies were recorded by scanning the entire volume (~20 slices with a step size of 1.5 μm) of the dorsal organ ganglion (**Movie 1**), while ORN axon terminals were recorded from a single slice of the antennal lobe (**Movie 2**). Dose-response experiments (data shown in **Fig 1-2**, **Supp Fig 3-4** and **Movie 1**) were performed using larvae of the *Orco>GCaMP6m*, *Orco>mCherry.NLS* genotype and recording from ORN cell bodies. White noise experiments (data shown in **Fig 4**, **Supp Fig 7-8** and **Movie 2**) were performed using larvae expressing GCaMP6m in a single ORN (e.g. *Or42a>GCaMP6m* used in **Supp Fig 7**) and recording from ORN axon terminals.

### Dose response analysis

Custom code written in ImageJ was used to track and identify each ORN as well as its responses to odorant stimuli. Slight movement artifacts were corrected by aligning frames using mCherry NLS labeling of ORN cell bodies and the ImageJ TurboReg plugin (Thevenaz et al, 1998). Each ORN activated in response to an odorant stimulus was visually identified using both the anatomical location of its dendritic bundle and the functional map of cognate odorant to ORN activation (**Fig 1 E, F**, **Supp Fig 2**). ORN identification was performed independently by two experimenters to ensure accuracy. Changes in fluorescence were then quantified as (*F*_Peak_ - *F*_0_)/*F*_0_, where *F*_0_ was the average ORN intensity sampled from the frames immediately preceding odorant delivery and *F*_peak_ was the highest intensity in ORN fluorescence during odorant delivery. Each odorant stimulus was repeated with at least 5 trials. The raw response data is summarized in **Supp Table 1**.

The heatmap in **Fig 2A** was generated by directly averaging the peak responses across trials. Simulated annealing was used to optimize the order of ORNs and odorants presented in this heatmap, such that it minimized a loss function in which cost increased linearly with the distance that activated odorant-ORN pairs were from the matrix diagonal. The response data was normalized by the highest response level within each trial, averaged across trials, and then Z-scored prior to performing PCA. The distance and direction of vectors shown in **Supp Fig 5** were calculated for each data point in **Fig 2B** using the standard formulae for cartesian to polar coordinate transformation.

### Dose-response curve fitting

A Hill equation with a unique set of parameter values was fit to the dose-response curve for each odorant-ORN pair in our data set. The general form of the Hill equation is as follows:

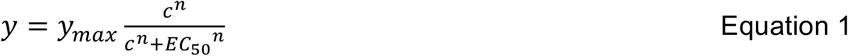

where *y*_*max*_ is the maximum ORN response level across concentrations, *c* is the odorant concentration, *EC*_50_ is the half-maximal effective concentration, and *n* is the Hill coefficient. In the calcium imaging experiment, the maximum fluorescence intensity *y*_*max*_ could be affected by the detailed experimental settings and it is differently fitted for the curve of each odorant-ORN pair. Here, the absolute value of *y*_*max*_ is not considered a coding feature, so in the following analysis, we normalized the responses using *y*_*max*_.

There were 21 odorant-ORN pairs saturated within the concentration range we studied. We started by fitting these 21 curves using the Hill equation. We normalized the responses using the *y*_*max*_ for each odorant-ORN pair and shifted the x-axis using its *EC*_50_. A scatter plot of the normalized and shifted dose-response data for the 21 odorant-ORN pairs is shown in **Fig 3A**.

Next, we used a Hill equation function with *y*_*max*_ = 1 and *EC*_50_ = 0 to fit all 105 data points from the 21 different odorant-ORN pairs. The resulting equation had a Hill coefficient n = 1.45, with R^2^ > 0.99. Next, we applied this Hill coefficient to fit odorant-ORN data pairs that did not saturate in the concentration range we had tested. There were 19 additional odorant-ORN pairs that were close to saturation and we could therefore estimate their *y*_*max*_ and *EC*_50_ values well.

After fitting the 21 odorant-ORN pairs that had saturated as well as the 19 that were close to saturation, we had at least one parametrized Hill equation for each odorant. To fit the remaining odorant-ORN pairs that were not close to saturation within our tested concentration range, we first assumed that each odorant had approximately the same *y*_*max*_ for each odorant (this was calculated by averaging the *y*_*max*_ of all ORNs that shared an odorant with a parameterized logistic curve). Given the known *y*_*max*_ for each odorant and the fixed Hill coefficient, we could estimate the *EC*_50_ for the remaining 100 weakly responding odorant-ORN pairs.

The *EC*_50_ of all odorant-ORN pairs is summarized in **Fig 3B**. The black elements in the matrix indicate that the corresponding ORN showed no activity within the tested concentration range; we were unable to fit an *EC*_50_ value for these odorant-ORN pairs. We used the Hill equation and fitted parameters for each odorant-ORN pair to generate the activity response data and found that it was similar to the actual data (**Supp Fig 6A**).

### Analysis of the *EC*_50_ Matrix

To perform PCA on the *EC*_50_ matrix, we first transformed the values to the *-ln*(*EC*_50_), such that odorant-ORN pairs with a high sensitivity (small *EC*_50_) were now represented by large values and those that were less sensitive (large *EC*_50_) had small values. The remaining pairs that that did not have an *EC*_50_value (the missing data, represented by black squares in **Fig 3B**), represent pairs with a much lower sensitivity and were set to zero. **Supp Fig 6B** shows the percentage of variance explained by each principal component (PC) once PCA was performed on the-ln (*EC*_50_) matrix. In comparison to a shuffled matrix (in which each row is randomly permuted), we found that only the first PC was significantly different (p < 0.0001 for 1000 instances of shuffled data).

We compared 32 descriptors of molecular structure from the E-Dragon software, which were found in Haddad et al., 2008 to be relevant for olfactory coding across animals. We found that one metric, aromaticity index of a molecule, had the highest correlation with the first PC of the *EC*_50_ matrix with a coefficient of 0.8 (**Supp Fig 6C**).

We fit the power law distribution using code from (Clauset et al., 2009). The resulting fitting index of 0.22 (large values mean better fit to the power law for this metric) is larger than the threshold (0.1) needed to accept the power law hypothesis (Clauset et al., 2009).

### Derivation of power law scaling of ORN ensemble responses from *EC*_50_ distribution

Here, we explain analytically the power law relation between odorant concentration and the ensemble response of ORNs. Under the same Hill equation we used to fit individual dose-response curves (Eq. 1, here we set **y*_*max*_ = 1 for simplicity), assume that (i) *EC*_50_ follows a power law distribution *P*(1/*EC*_50_) ∝ (1/*EC*_50_)^-λ-1^ (or equivalently an exponential distribution for *k* =-ln (*EC*_50_): *P*(*k*) = λe^-λ(*k*-*k*_0_)^,*k* ≥ *k*_0_)* (ii) the Hill coefficient *n* for all odorant-ORN pairs are the same and greater than *λ* (satisfied in the data as 1.45 vs. 0.35). If so, the ensemble response follows an approximate power law form *r*(*c*) ∝ *c*^*λ*^ for concentrations *c* ≤ *e*^-*k*_0_^ (which means the weakest response pair in the ensemble has not reached the half level). For convenience, we use the log scale of concentration and *EC*_50_: *x* = ln(*c*), *k* =-ln (*EC*_50_) and the logistic function in place of the Hill equation: 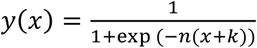.

This result can be intuitively obtained by considering the limiting case where the logistic function is infinitely steep (large Hill coefficient) and is thus replaced by a step function. The ensemble response combining a large number of odorant-ORN pairs can be expressed as an integral: 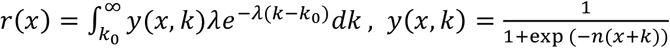 is the log-concentration. When *y*(*x,k*) is a step function, the integral becomes *P*(*k* ≥ -*x*), which is essentially the cumulative density function for *k*. Given the distribution of *k*, this is exactly an exponential function *r*(*x*) = *e*^*λ*(*x* + *k*_0_)^ (or a power law function of *c*) for *x* ≤ -*k*_0_, and saturates at larger concentrations.

For the general case of logistic activation, the integral does not have a simple form expression but involves hyper-geometric functions. However, we can derive a simple closed form approximation by approximating the logistic function *f*(*x*) = 1/(1 + *e*^-*nx*^) using piecewise exponential functions:

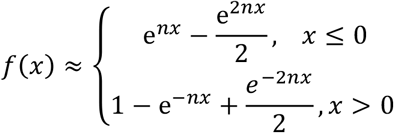

Such an approximation becomes asymptotically exact when the steepness *n* goes to infinity, or when the absolute value of x goes to infinity. Substituting *y*(*x,k*) with this approximation, the integral splits into segments, over which the integrand are sums of exponential functions, and therefore can be easily integrated. This gives the closed form approximation of *r*(*x*):

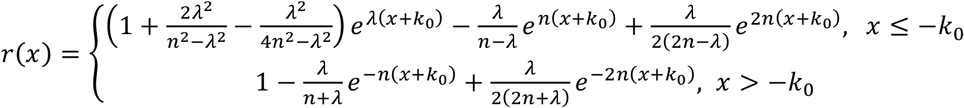

For small concentrations, *x* ≤ -*k*_0_, the leading term in the above expression is *e*^λ(*x* + *k*_0_)^, since *λ* < *n*. This explains that the ensemble response is approximated by an exponential function with exponent *λ*. Furthermore, the theory also predicts the magnitude (vertical shift in the log-log plot of ensemble response as in **Fig 3C**), that is, 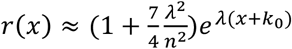, which explains how the Hill coefficient affect the ensemble response.

### Reverse-correlation analysis

White noise experiments were performed in a manner similar to those described in Kato, et.al, (2014). Briefly, we used custom code written in MATLAB to control odorant and water switching such that it followed an m-sequence. Calcium imaging was performed on the axon terminal of individual ORNs at ~30 frames per second. Calibration and an example of such a recording is shown in **Movie 2**. We then used a linear-nonlinear model to compare the m-sequence input to ORN responses during a 150second interval (from 60 - 210 sec). An 18 second time window was used for the linear filter, of which 15 seconds represented stimulus history in order to ensure extraction of the full filter dynamics (**Supp Fig 7B**). Next, we applied the linear filter to the data and compared this to the output in order to capture the nonlinear function. We found that a sigmoidal function fits the nonlinear function well (**Supp Fig 7C**). We applied novel m-sequences to validate the linear-nonlinear model (**Supp Fig 7D**) and found that they fit the data well. Peak and decay times for each filter were found by extracting the time points corresponding to the maximum amplitude and half maximum amplitude of the decay phase, respectively. 454 filters were calculated from the recording of 138 larvae responding to various m-sequence stimuli. Each of the 31 filters quantified in **Supp Fig 7**, are averaged across 10 trials.

Data, code and software can be found at: <u>https://github.com/samuellab/larvalolfaction</u> Microfluidic device pattern design can be found at: <u>https://metafluidics.org/devices/larvalolfaction/</u>.

**Supplementary Figure 1.**
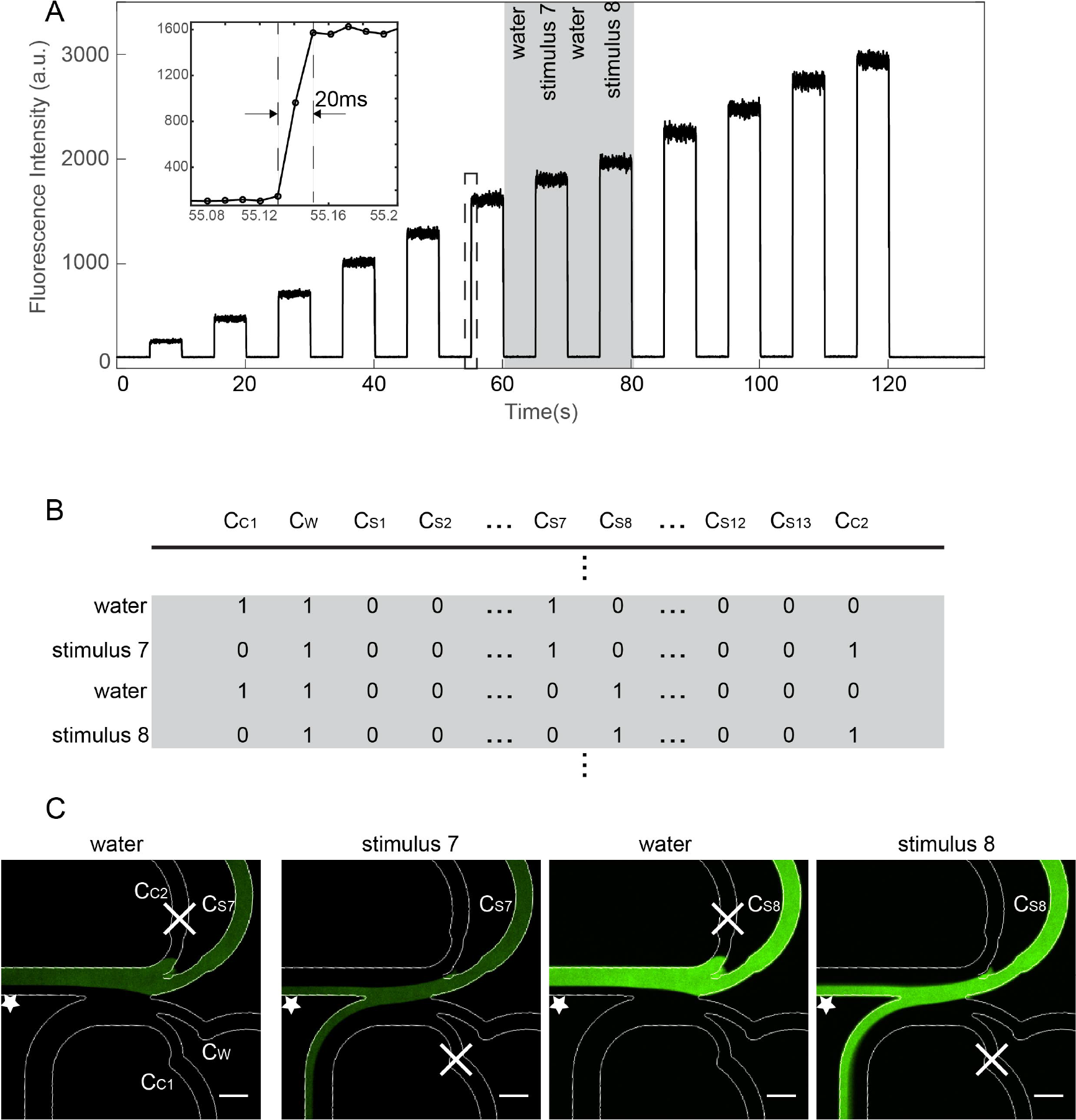
Validation of temporal and spatial odorant profiles in microfluidics device, using a fluorescent dye. A. Change in fluorescence intensity during delivery of 5 sec step pulses of increasing concentration of fluorescein dye, each followed by 5 second of water. Inset shows zoom-in of dashed box, indicating stimulus transition time is ~20 ms. B. Combination of valve states required to generate the stimulation sequence in shaded area of panel A; 1 and 0 indicate valve is open or closed, respectively. C_W_ represents water channel, C_C1_ and C_C2_ represent control channels that allow stimulus switching, and C_7_ and C_8_ represent odorant delivery channels and only open prior to and during stimulus delivery. C. Images of fluorescein dye, representing an odorant stimulus, in the microfluidics device during each state shown in panel B (water, stimulus 7, stimulus 8). Cross mark indicates closed channels, star marks the location of the larva’s “nose”. Scale bar is 300 μm.

**Supplementary Figure 2.**
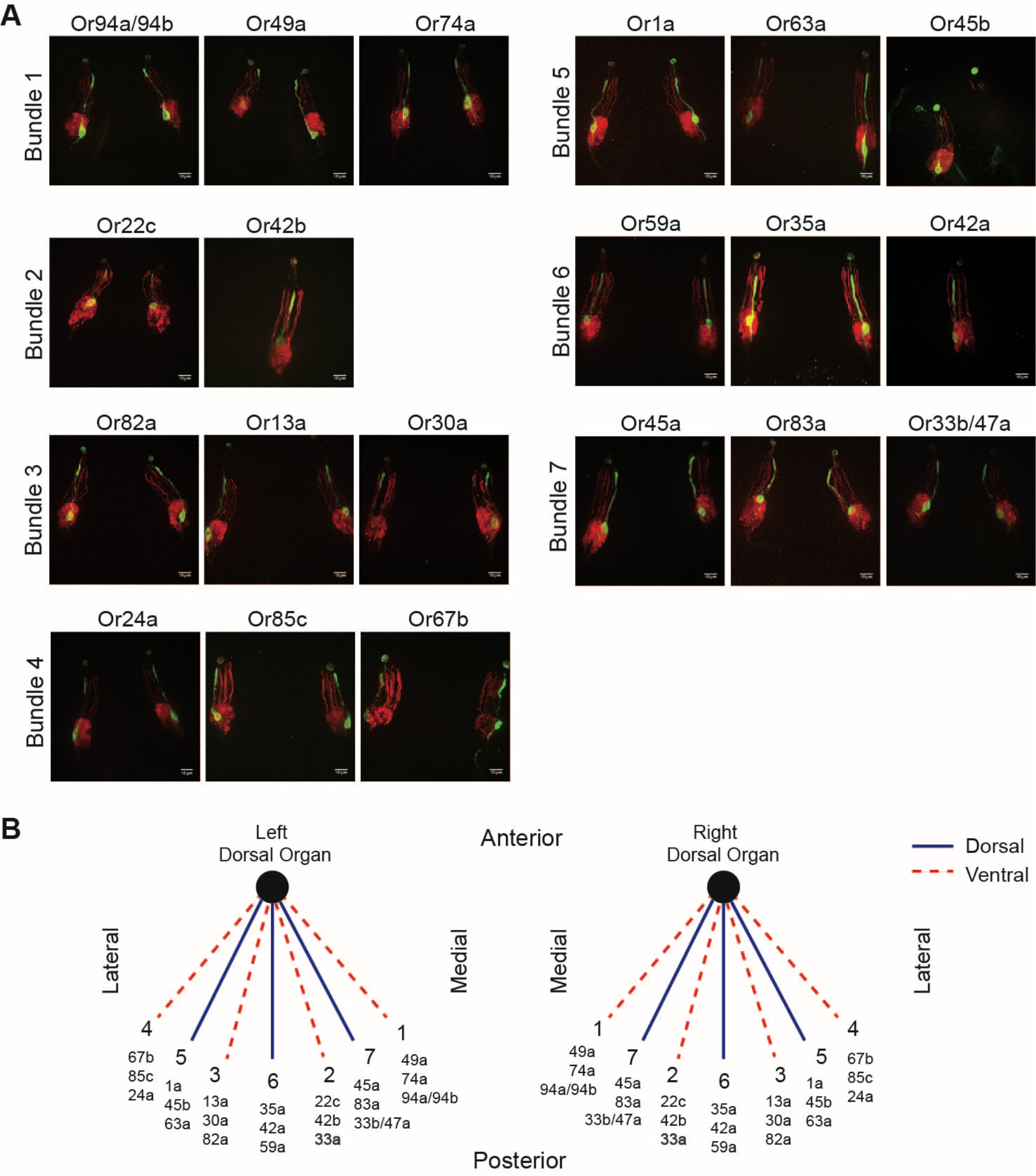
Anatomical map of ORN dendritic organization. A. Image of each ORN’s dendritic location using GFP to label a specific ORN and RFP to label all ORNs. Larvae expressing *OrX>GFP*, *Orco>RFP*, where OrX is a specific olfactory receptor. We infer the vacancy in bundle 2 is Or33a. During functional imaging, there were no strong signals from this neuron. No expression of Or2a and Or7a were observed in first instar larvae. B. Summary schematic of stereotyped ORN position in each dendritic bundle for left and right dorsal organs.

**Supplementary Figure 3.**
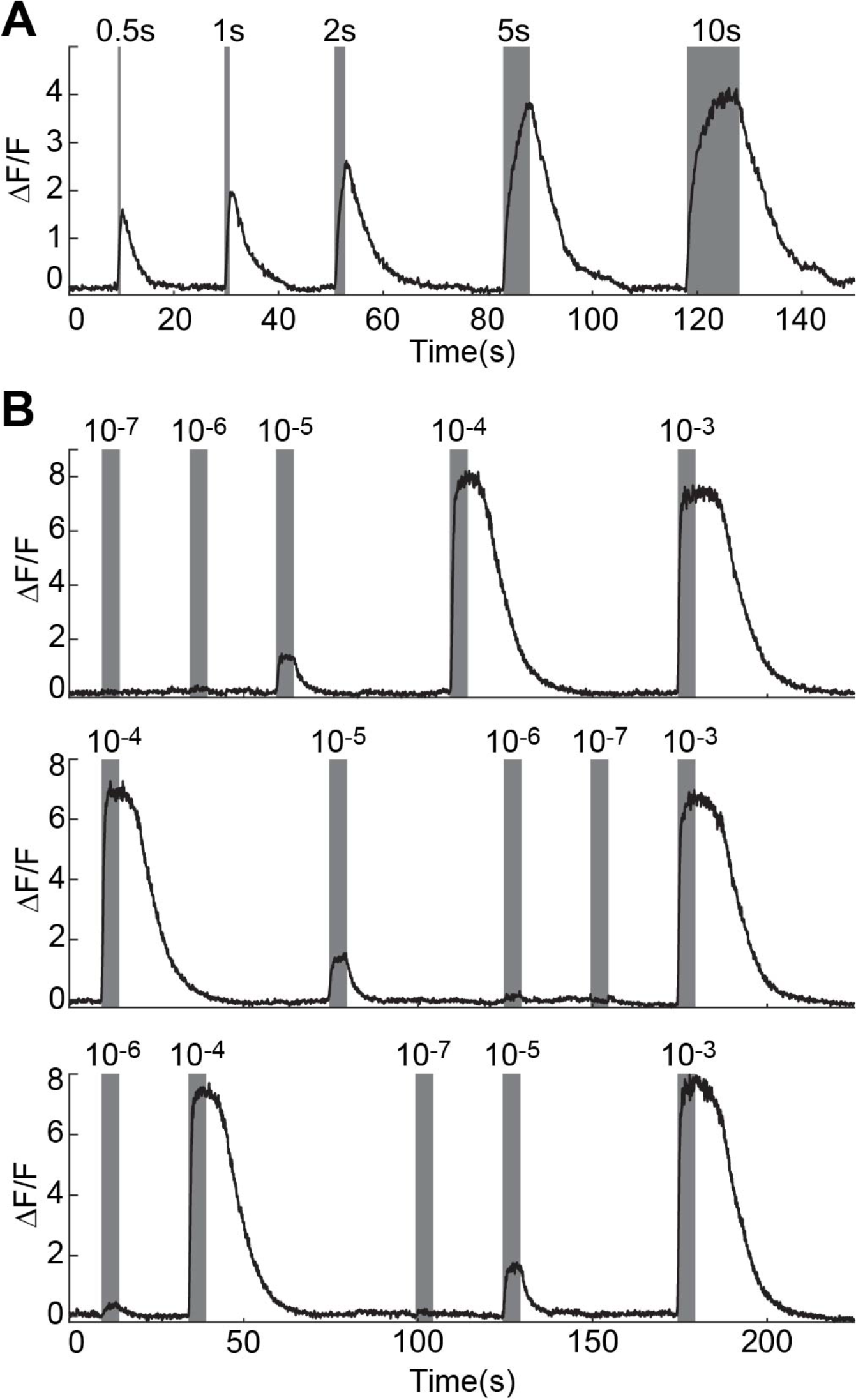
Effect of stimulus duration and sequence on ORN response. A. Or35a-ORN responses to 0.5, 1, 2, 5 and 10 seconds of 10^-5^ dilution of 3-octanol. The maximum response saturates when odorant pulse is longer than 5 seconds. B. Or35a-ORN response to increasing (top panel), primarily decreasing (middle panel), and random (bottom panel) concentration sequences of 3-octanol pulses, delivered at 5 seconds each. The response amplitude to each concentration level is history independent.

**Supplementary Figure 4.**
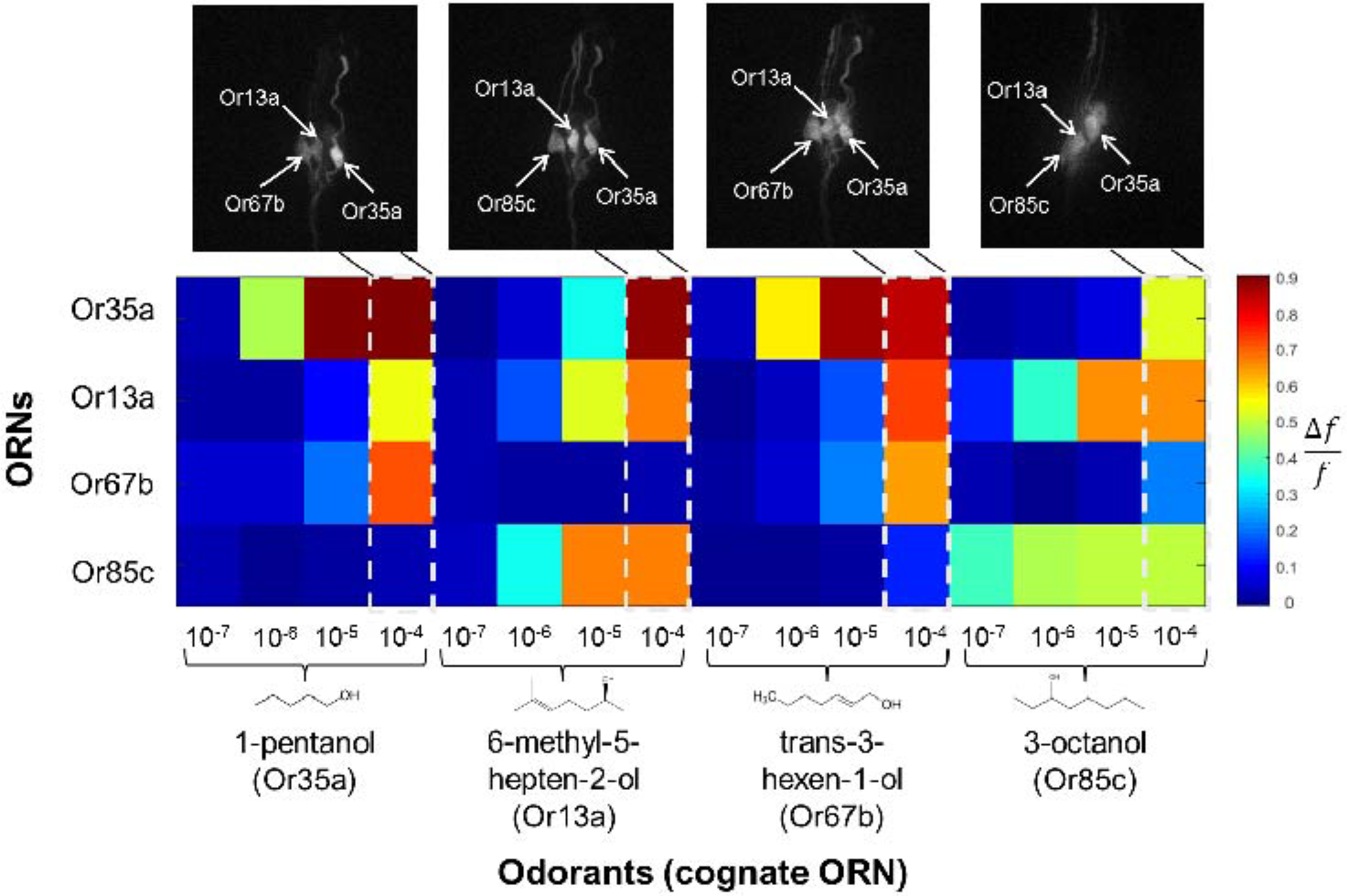
Coactivation of ORNs with similar cognate odorants at high concentrations. Heatmap of peak responses of four ORNs to four alcohol odorants, across four concentrations of each odorant. Activities normalized by maximum response amplitude. Neural images show responsive ORNs in dorsal organ ganglion during calcium imaging at the highest odorant concentrations.

**Supplementary Figure 5.**
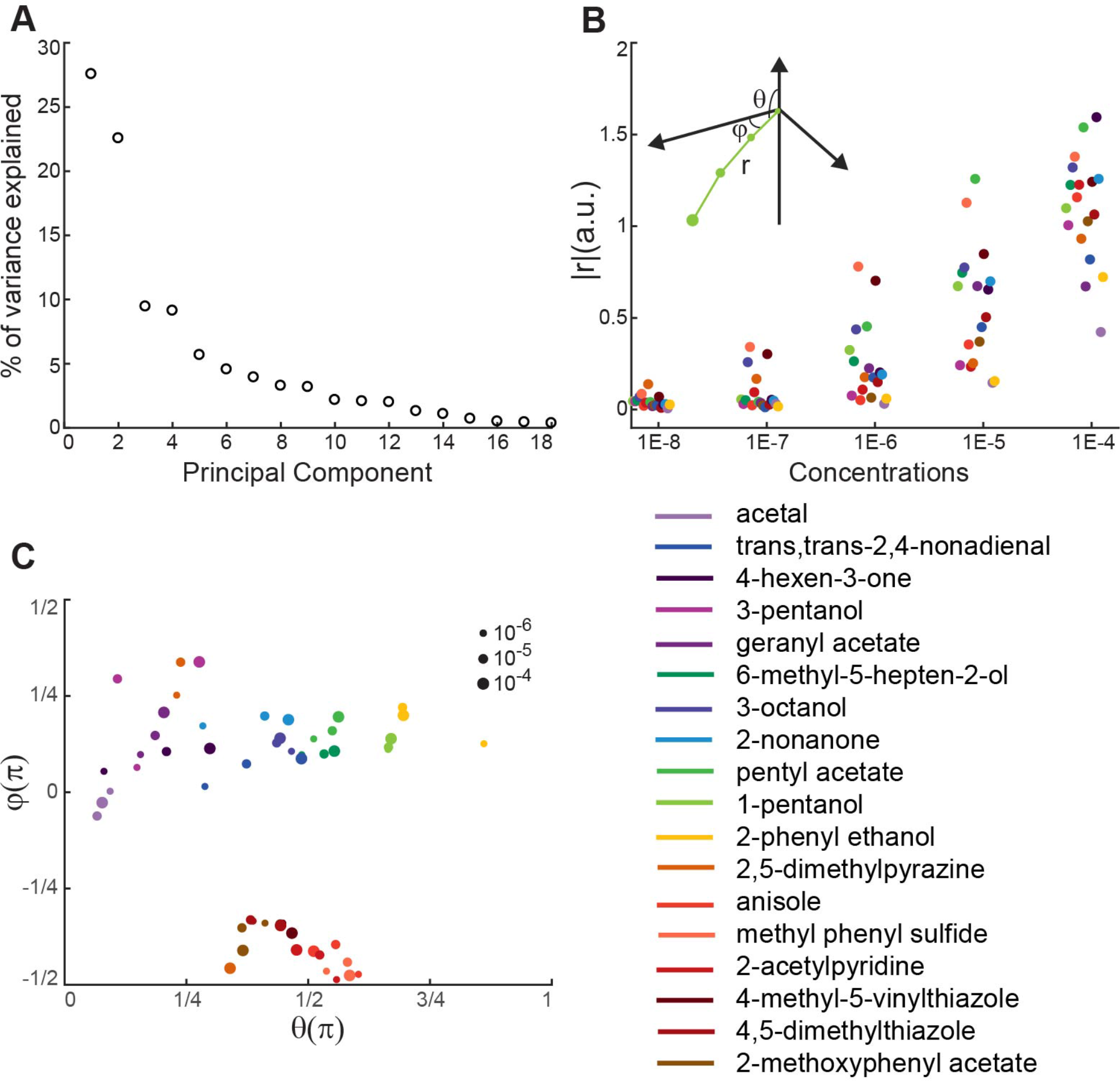
PCA analysis of ORN dose-response data. A. Percentage of variance explained by each principal component. B. and C. Transform of odorant vectors in PCA space (**Fig 2B**) to spherical coordinates (inset of B). Odorant vector length increases monotonically with increasing concentration. Angular direction of different odorants (represented by dot color) separate, but aggregate for direction of different concentrations of the same odorant (dot size).

**Supplementary Figure 6.**
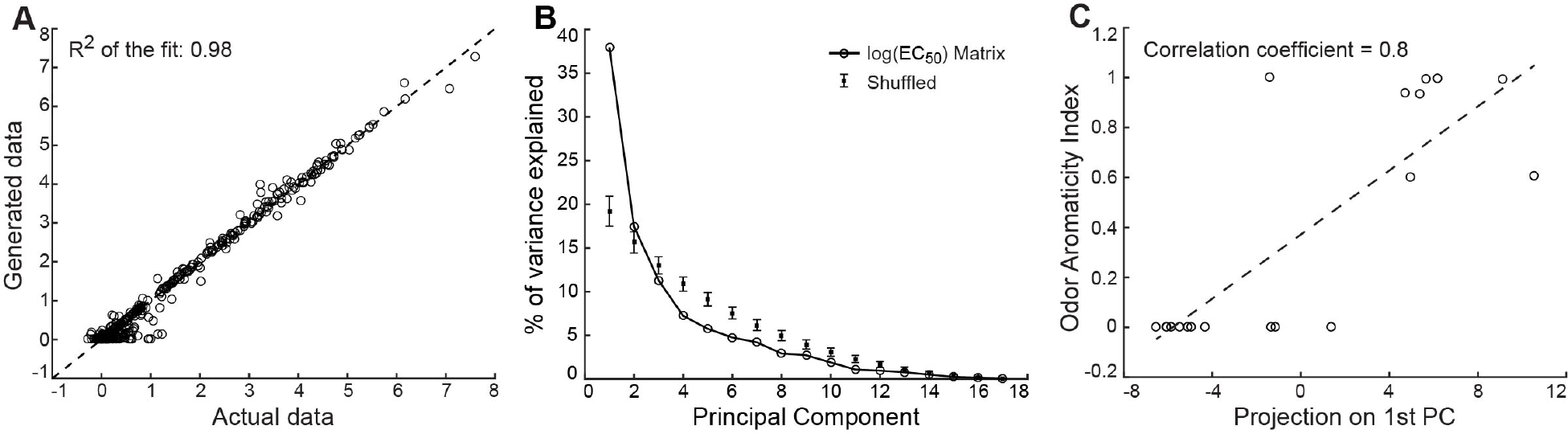
Aromaticity index highly correlated to major PC of EC_50_ matrix. A. Actual neural activity data is highly correlated with simulated data generated using the Hill equation and fitted parameters. Scatter plot includes all non-zero data in **Fig 2A**. Dashed line indicates *y* = *x.R*^2^ is 0.98. B. Percentage of variance explained by each PC of the PCA on the-ln (*EC*_10_). Data compared with the corresponding results from 1000 randomly shuffled data. C. Correlation plot between each odorants projection on the 1^st^ PC of-ln (*EC*_10_) matrix and its aromaticity index.

**Supplementary Figure 7.**
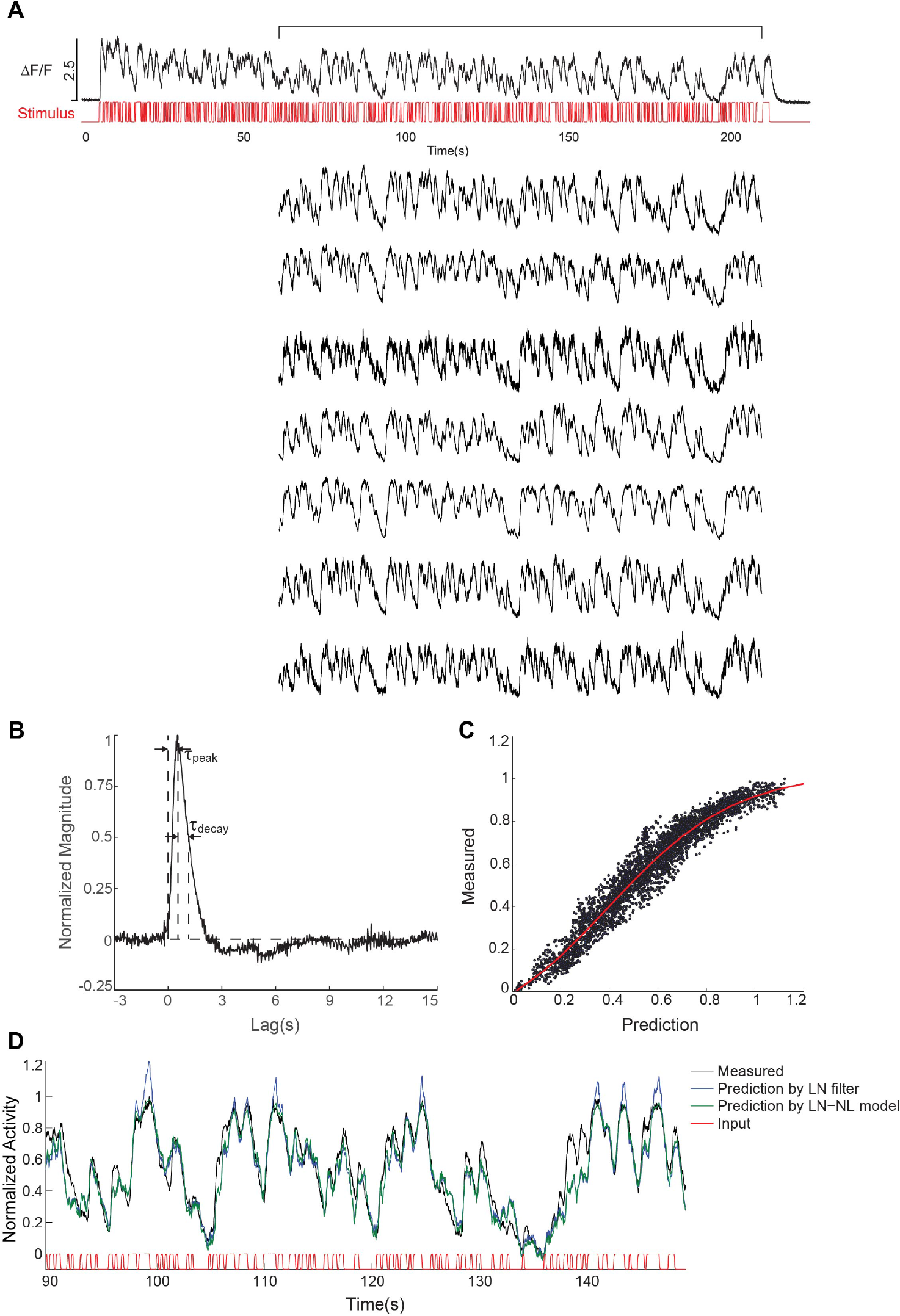
Linear-nonlinear model of ORN temporal dynamics. A. Multiple trial responses of Or42a-ORN to the same m-sequence of 10^-7^ dilution of 3-pentanol. B. The linear filter calculated from reverse correlation of the input-output from the first trial of panel A. C. Non-linear transfer function calculated by comparing measured and predicted responses using the linear filter. D. Validation of the linear-nonlinear model by comparing predicted and measured responses to novel m-sequence stimuli.

**Supplementary Figure 8.**
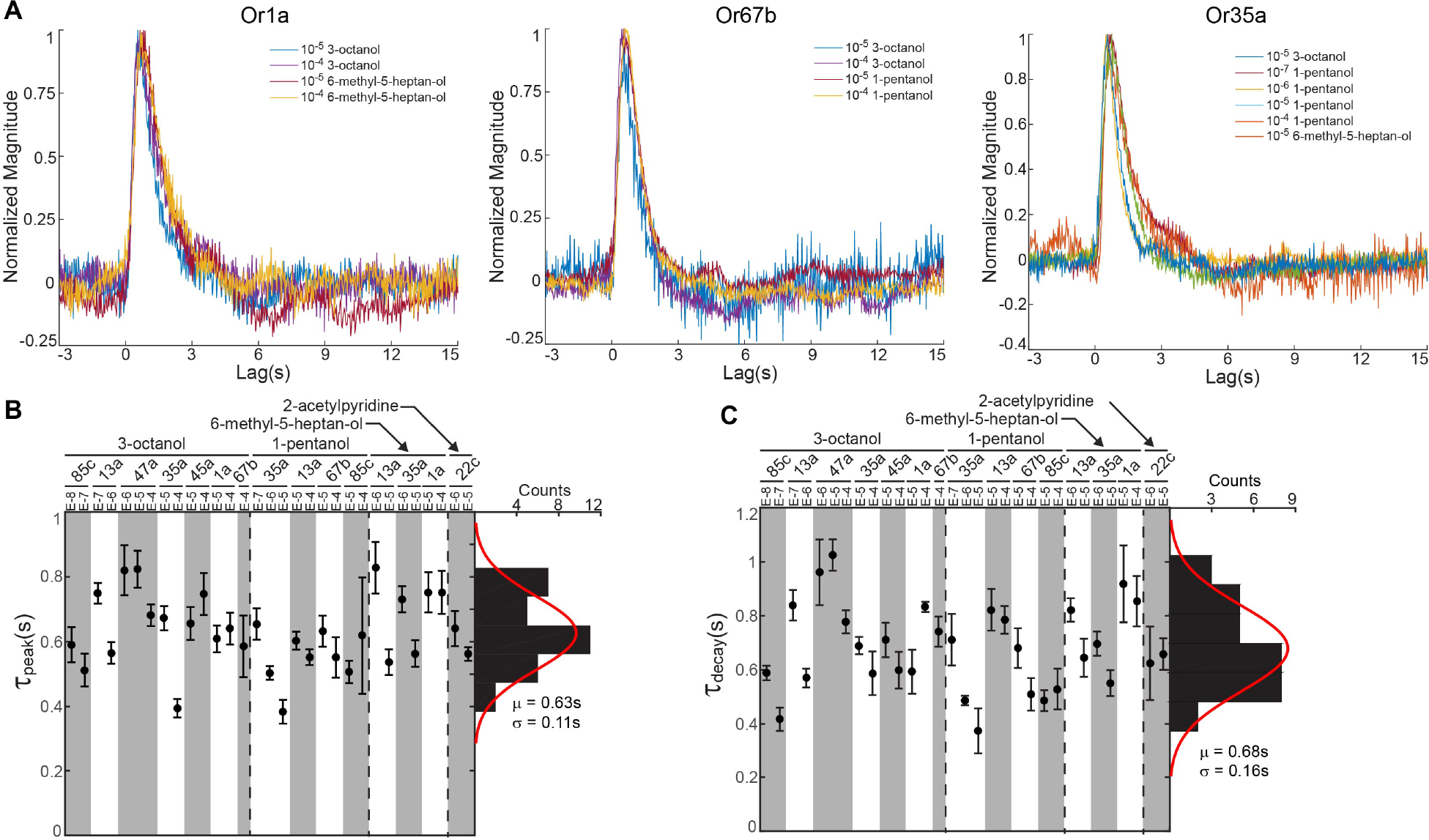
Comparison of ORN temporal filters. A. Normalized filters of three ORNs responding to various odorant stimuli. B. and C. Distribution of peak time (B.) and decay time (C.) of 31 filters measured from various ORN and odorant stimuli. Distributions of peak and decay times were fit to Gaussian distributions with mean and variance labeled below histogram.

**Movie 1. Dose-dependent activation of ORNs**. Left: Calcium imaging of 21 pairs of larval ORNs in response to increasing concentrations of the 1-pentanol odorant, from 10“^7^ to 10“^3^ dilutions. Movie starts by scanning through the imaging volume to identify ORNs activated at the highest concentration level. Three ORNs are responsive on the left side and five on the right side. ORN identity was confirmed from the dendrite location and response to panel of 13 cognate odorants (not shown in the movie). Right: Responses of ORNs to step pulses of odorant stimuli.

**Movie 2. ORN responses to pseudorandom white noise stimulus**. Top left, stimulus delivery marked by fluorescence. Bottom left, axon terminal of Or45a-ORN responding to the white noise stimulus using 10^-7^ dilution of 2-nonanone. Top and bottom right, real time plots of the input stimulus and ORN response during an experiment.

**Supplementary Table 1**. Raw activity data of 21 ORNs responding to 19 odorants at five concentration levels collected from 122 recordings.

